# Site-specific glycosylation mapping of Fc gamma receptor IIIb from neutrophils of health donors

**DOI:** 10.1101/2020.04.29.068742

**Authors:** Iwona Wojcik, Thomas Sénard, Erik L de Graaf, George MC Janssen, Arnoud H de Ru, Yassene Mohammed, Peter A van Veelen, Gestur Vidarsson, Manfred Wuhrer, David Falck

**Author notes:** Both authors contributed equally (Iwona Wojcik, Thomas Sénard).

## Abstract

Fc gamma receptors (FcγR) translate antigen-recognition by immunoglobulin G (IgG) into various immune responses. A better understanding of this key element of immunity promises novel insights into mechanisms of (auto-/allo-)immune diseases and more rationally designed antibody-based drugs. Glycosylation on both IgG and FcγR impacts their interaction dramatically. In this study, we developed a straightforward and comprehensive analytical methodology to map FcγRIIIb glycosylation from primary human material. In contrast to recently published alternatives, we assessed all glycosylation sites in a single LC-MS/MS run and simultaneously determined the donor allotype. Studying FcγRIIIb derived from healthy donor neutrophils, we observed profound differences as compared to the soluble variant and the homologous FcγRIIIa on natural killer cells. This method will allow assessment of FcγRIII glycosylation differences between individuals, cell types, subcellular locations and pathophysiological conditions.

## INTRODUCTION

Binding of immunoglobulin G (IgG) to the Fc gamma receptors (FcγR) initiates and regulates immune cell signalling which results in important immune responses^1,2^. Therefore, FcγR has a key role in homeostasis and in many pathologies^3,4^. This is also widely exploited for therapeutic purposes, for example with monoclonal antibodies or polyclonal intravenous IgG^5-7^. The key interaction between IgG and FcγR is heavily regulated by the proteoform distribution of either binding partner, e.g. through post-translational modifications. The impact of IgG proteoforms has been extensively studied in the last decades, though a complete picture still eludes us^8,9^. Subclass, allotype and glycosylation, especially (a)fucosylation, of IgG impact FcγR binding^10-12^. IgGs convey their effector functions for a large part via Fcγ receptors^13^. These effector functions are regulated by their varying expression on different immune cells. For example, FcγRIIIa is an activating FcγR found on natural killer (NK) cells, monocytes and macrophages. Like other activating receptors it can mediate antibody-dependent cellular cytotoxicity (ADCC) or phagocytosis, respectively through the associated signalling γ-chain containing intracellular Tyrosine-based Activation Motif (ITAM). On the other hand, FcγRIIb down-regulates inflammatory responses through its inhibitory action on several immune cells through its Immunoreceptor Tyrosine-based Inhibition Motif (ITIM)^1^. Unlike all other FcγR, the human FcγRIIIb is GPI-anchored and lacks a transmembrane domain. It is found expressed mainly on granulocytes and seems to affect signalling of other Fc-receptors through accumulation in lipid rafts enriched in kinases (Src) required for ITAM-phosphorylation and signalling^14^. Given the relatively high expression levels of FcγRIIIb on neutrophils with 200,000 to 300,000 copies per cell^15^, and the dominance of neutrophils amongst white blood cells, FcγRIIIb can be considered the most abundant FcγR in circulation. Known functions include activation of neutrophil degranulation, cell adhesion, calcium mobilization and neutrophil tethering to soluble immune complexes^16-19^.

Despite recent advances, the role of FcγR proteoforms is only poorly understood^20^. For FcγR, allotypes lead to differentially active proteoforms. The known allotypes of FcγRIIIb, neutrophil antigen 1 (NA1) and 2 (NA2), and SH (SH being rather uncommon), have been shown to differ in their affinity for IgG and capacity to induce phagocytosis of IgG opsonized targets^21^. Glycosylation of FcγR can also strongly impact the interaction with IgG. Some receptor glycans have direct glycan-glycan and glycan-protein interactions with bound IgG^22,23^. Deglycosylation of site N_162_ (unique to FcγRIIIa and FcγRIIIb), strongly elevates affinity of FcγRIIIa to IgG, but also alleviates sensitivity to IgG-core-fucosylation^24,25^. Furthermore, differences in FcγRIIIb glycosylation in different cell models, have been shown to impact IgG binding^26,27^. The available studies underline the importance of FcγR glycosylation, but can so far only sketch a very rough picture of its functional impact.

A prominent reason for this lack of functional understanding is the limited availability of data on FcγR glycosylation of primary human material^20^. While the great heterogeneity of proteoforms, especially in FcγRIIIb, was already apparent in early studies^15^, glycomics studies on primary human material only became possible in recent years^28^. However, owing to the great complexity and differential functional impact of glycosylation sites, only site-specific glycoproteomics studies can characterize FcγR glycosylation to the necessary extent. These recent glycoproteomics studies focused on healthy volunteers. They revealed FcγRIIIa glycosylation of NK cells^29^ and monocytes^30^ and FcγRIIIb glycosylation of neutrophils^31^. The basis was purification of the cells by negative selection with magnetic beads, followed by immunoprecipitation of FcγR. Additionally, the soluble FcγRIIIb^32^, originating from shedding from neutrophils upon activation, has been purified and studied from plasma. The purified receptor from all sources was analysed by bottom-up glycoproteomics following protease cleavage with chymotrypsin and/or endoprotease GluC^33^.

While ground-breaking, the two previous studies on site-specific glycosylation of FcγRIIIb also had some limitations^31,32^. Both studies relied on two independent proteolytic cleavages to assess different sites, thus necessitating multiple liquid chromatography – mass spectrometry (LC–MS) runs to cover the whole receptor. Yagi *et al*. focused on the soluble version of FcγRIIIb which is released from neutrophils during activation and whose function is largely unknown^32^. They used pooled blood from multiple donors, loosing inter-donor variability. Washburn *et al*. only analysed three of the six potential *N*-glycosylation sites, but of 50 donors accumulating strong data on inter-donor variability^31^. Nonetheless, we still know too little about the functional and clinical impact of FcγR glycosylation to prefer a method focussing only on certain glycosylation sites. Other existing strategies covering all sites of FcγRIIIb or FcγRIIIa are quite complicated and difficult to apply to clinical investigations where eventually large numbers of samples need to be detected in a robust way^29,32^.

Here, we present a method for the site-specific analysis of all glycosylation sites of FcγRIIIb in a single LC-MS/MS experiment. With it, we identified and relatively quantified neutrophil-derived FcγRIIIb glycosylation individually for multiple donors. Additionally, our approach allowed a qualitative overview of site occupancy and the determination of donor allotypes. This was enabled by avoiding glycopeptide enrichment which also promises more robustness. Additionally, interferences from endogenous IgG and from leaking capturing antibody are avoided by a simple non-reducing SDS-PAGE step. Additionally, our method is generic for FcγRIIIa and FcγRIIIb, making it potentially applicable to a wide range of leukocytes. Moving towards clinical investigations of FcγR glycosylation will be essential for a complete understanding of the (patho-)physiological role of IgG-FcγR interactions. Our methodology presents a uniquely suited starting point, as it is unprecedented in its ability to simultaneously cover individual donor, subclass, allotype, cell and site differences of FcγRIII glycosylation comprehensively.

## RESULTS AND DISCUSSION

### FcγRIIIb purification and identification

FcγRIII was immunoprecipitated from ∼16 million primary human neutrophils cells from healthy donors (**Supplementary Table S1**). FcγRIIIb is known to be very abundant on human neutrophils, while FcγRIIIa is not present^19,34^. However the absence of FcγRIIIa from neutrophils has recently been challenged^35^. The western blot showed a smear from 50 to 80 kDa when probed with an anti-CD16 antibody (**Figure 1**). This behaviour, previously reported for example by Galon *et. al* ^36^, confirmed the presence of FcγRIIIb with its abundant and diverse glycosylation pattern. The non-reducing SDS-PAGE purification aided in separation of FcγRIIIb from interferences derived, for example, from the capturing antibody or endogenous IgG (**Supplementary Figure S1**). This is simple and preferable to an affinity removal of IgG, e.g. with protein G, which may lose specific FcγR proteoforms due to high affinity interactions^29^. A comparison of total cell lysate, flow-through and eluate demonstrates the efficacy of the purification (**Figure 1, Supplementary Figure S1**). Furthermore, MS/MS analysis of the purified protein yielded a sequence coverage of approximately 80% and did not indicate any major interferences in the relevant gel band (**Supplementary Table S2**).

**Figure 1.**
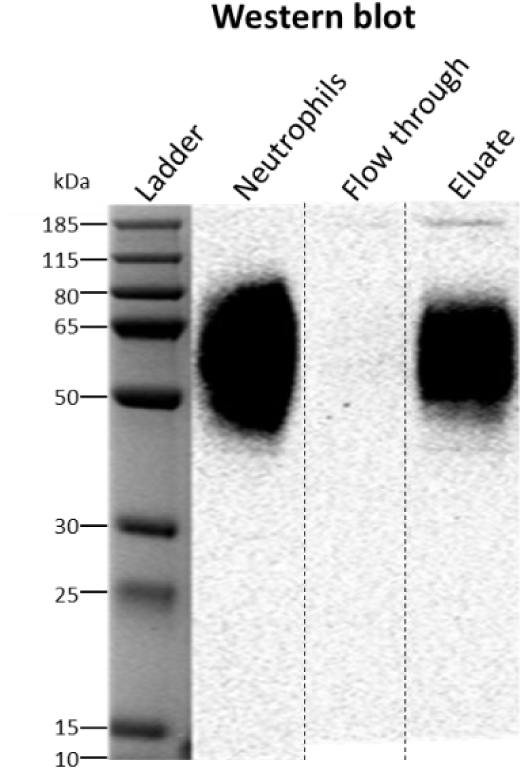
Western blot of human neutrophil lysate before and after FcγRIII immunoprecipitation. The identity of FcγRIIIb was confirmed by western blot. The flow-through lanes show the diluted *unbound* fraction of the immunoprecipitation while the eluate lanes present the purified FcγRIIIb protein. As expected, FcγRIIIb exhibits an elongated band from 50 to 80 kDa. Only part of the blot (**Supplementary Figure 1**) is shown. The cropping area is indicated by striated lines.

### Coverage and protein identity

Isolated FcγRIIIb was subjected to an in-gel endoproteinase GluC and chymotrypsin treatment prior the LC-MS(/MS) analysis^33^. This approach was sufficient to generate and identify unique peptides for all six glycosylation motifs. FH**N**_**45**_ESLISSQASSY (NES), FIDAATV**N**_**64**_DSGEY (NDS), RCQT**N**_**74**_LSTLSDPVQLE (NLS), CRGLVGSK**N**_**162**_VSSE (NVS) and TV**N**_**169**_ITITQGLAV (NIT) were found in human neutrophils (**Supplementary Table S3**), while SPED**N**_**38**_ESQW was only detected in recombinant material (data not shown). An overview of generated FcγRIIIb glycopeptides is depicted in **Figure 2**. The main difference between current and previous analytical attempts was that our method required only one LC-MS/MS run to obtain full coverage of all glycosylation sites, instead of separate proteolytic treatment and LC-MS/MS runs per site.

**Figure 2.**
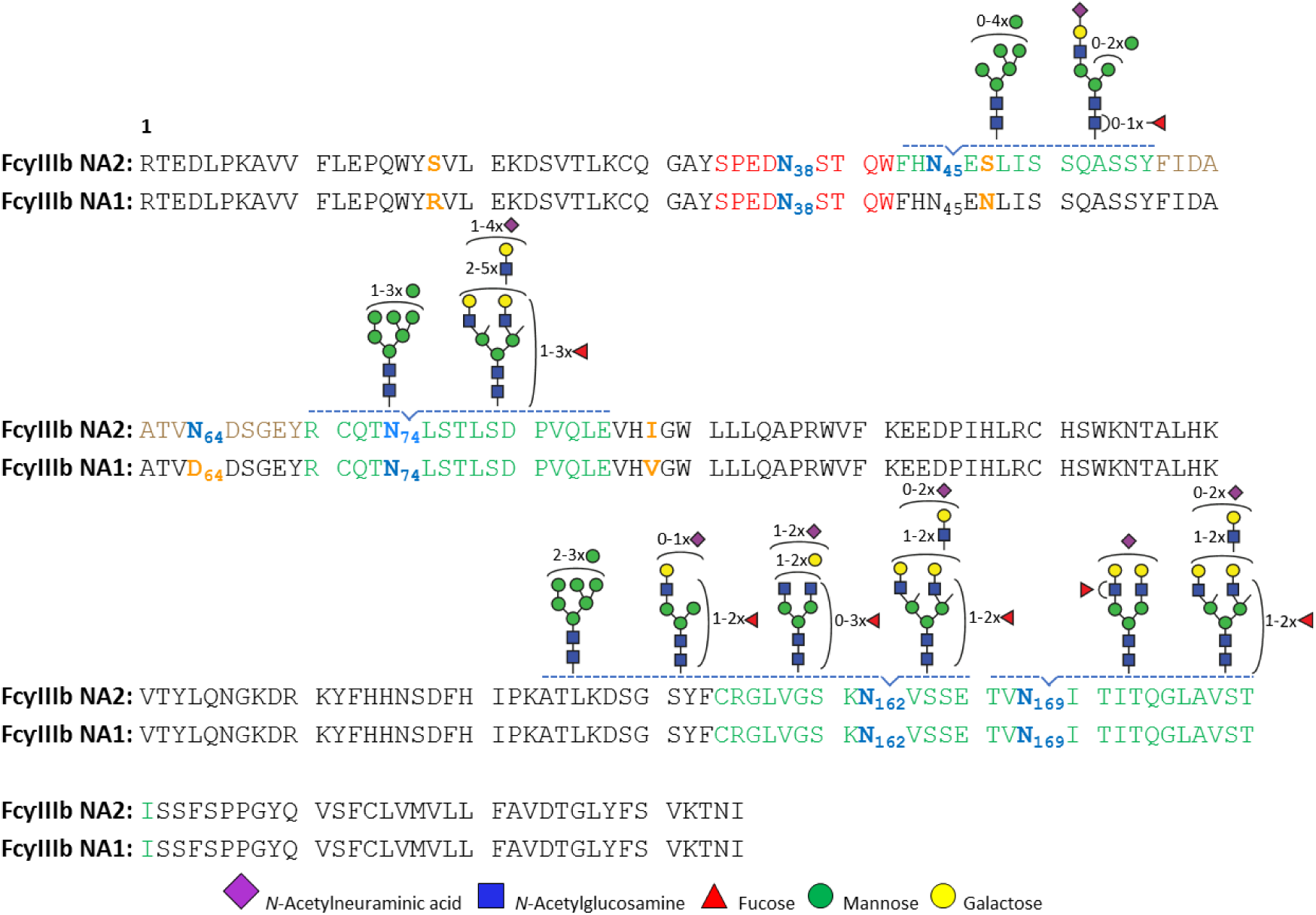
A schematic representation of site-specific N-glycosylation of endogenous FcγRIIIb NA1 and NA2 from human neutrophils. N-glycosylation sites are noted in bold, blue letters. All determined and occupied glycopeptides are depicted in green with their corresponding set of glycans. Peptides with an unoccupied N-glycosylation site (site N_45_ NA1 and N_64_ NA2) are marked in brown, while the sequence of a peptide containing site N_38_, that was only detected in recombinant material, is indicated in red. Amino acids in yellow show the sequence variations between NA1 and NA2.

The two major FcγRIIIb allotypes, NA1 and NA2 differ in four amino acids (see **Figure 2**), resulting in four and six potential glycosylation sites for NA1 (N_38_, N_74_, N_162_, N_169_) and NA2 (N_38_, N_45_, N_64_, N_74_, N_162_, N_169_), respectively^37,38^. Regarding functional differences, the NA2 allotype has been shown to possess a lower FcγR-mediated phagocytic capacity^21^. FcγRIIIb differed in electrophoretic mobility between the donors (**Supplementary Figure S2**). For Donor 1, FcγRIIIb showed a lower average molecular mass than for Donors 2 and 3 which may indicate the less glycosylated NA1 allotype. The FcγRIIIb allotype can be determined by proteomics^31^, which we achieved simultaneously to the glycoprofiling with our approach. The assignments were based on the presence and the intensity ratio of allele-specific peptides unique to NA1-or NA2-carrying individuals: for example, FHN_**45**_ENLISSQASSY (NEN), FIDAATVD_64_DSGEY (DDS) for NA1, and FH**N**_**45**_ESLISSQASSY (NES), FIDAATV**N**_**64**_DSGEY (NDS) specific for NA2 (**Supplementary Table S4**). Indeed, Donor 1 was an NA1/NA2 individual, with a 1:5 ratio of relative contribution of N_45_ oligomannose glycopeptides from NA2 (NES/(NES+NEN)). A bias in relative quantitation may be caused by reduced ionization efficiency of glycopeptides compared to non-glycosylated peptides. However, a similar ratio (1:4) for unoccupied N_64_ peptide (NDS/(DDS+ NDS)) was observed. Thus, the 1:5/1:4 ratio is likely to reflect the real expression levels between NA2 and NA1, which may also be influenced by naturally occurring gene copy number variation of FcγRIIIb^39,40^. Donor 2 and 3 were identified as NA2 homozygous individuals, since they contained approximately 98% of FH**N**_**45**_ESLISSQASSY and FIDAATV**N**_**64**_DSGEY peptides (**Supplementary Table S4**). The residual 2% were explained by shared peptides sequences between FcγRIIIb NA1 and FcγRIIIa (FHN_45_ENLISSQASSY, FIDAATVD_64_DSGEY and FIDAATVD_64_DSGEYR). We cannot exclude that these signals may be derived from contamination by NK cells, macrophages and/or monocytes. Additionally, deamidation of site N_64_ may contribute to the D_64_ peptide signals, but does not explain the N_45_ signals. Importantly, at these low levels, impact of FcγRIIIa glycopeptides on FcγRIIIb glycosylation profiling can be neglected. Therefore, co-isolation is preferred as it enables use of the same sample preparation protocol for FcγRIIIa dominated cell types.

We did not observe peptides or glycopeptides in the healthy donors corresponding to the glycosylation site N_38_. Of note, we were able to detect small glycans on recombinant FcγRIIIb (data not shown). Yagi *et al*. previously reported on site N_38_ large, highly branched glycan structures in the range of *m/z* 1400 to *m/z* 2000^32^.

### Glycopeptide identification

Profiling of *N*-glycosylation microheterogeneity resulted in the identification of 10 *N*-glycan compositions at Asn_45_, 15 at Asn_74_, 30 at Asn_162_ and 6 *N*-glycan compositions at Asn_169_, respectively. Of these 61, 36 glycoforms were confirmed by tandem mass spectrometry (**Supplementary Table S5**). Based on glycopeptide fragmentation data, mass accuracy and isotopologue pattern, together with knowledge on biosynthetic pathways, we propose the *N*-glycan structures shown in **Figure 3**. Numerous structural isomers could be present for the same *N*-glycan composition. We identified the presence of multiple isomeric structures, but with the current method we were not able to resolve them quantitatively. We confirmed the presence of sialic acid by the diagnostic ion at *m/z* 292.103. Antennary fucosylation was confirmed by presence of a B-ion at *m/z* 512.197 [hexose+*N*-acetylhexosamine+fucose+H]^+^. In contrast, core fucosylation was indicated by the formation of the ion assigned as [peptide+*N*-acetylhexosamine+fucose+H]^+^. *N*-glycans containing *N*-acetyllactosamine (LacNAc) repeats, were indicated by signals at *m/z* 731.272 [*N*-acetylhexosamine_2_+hexose_2_+H]^+^.

**Figure 3.**
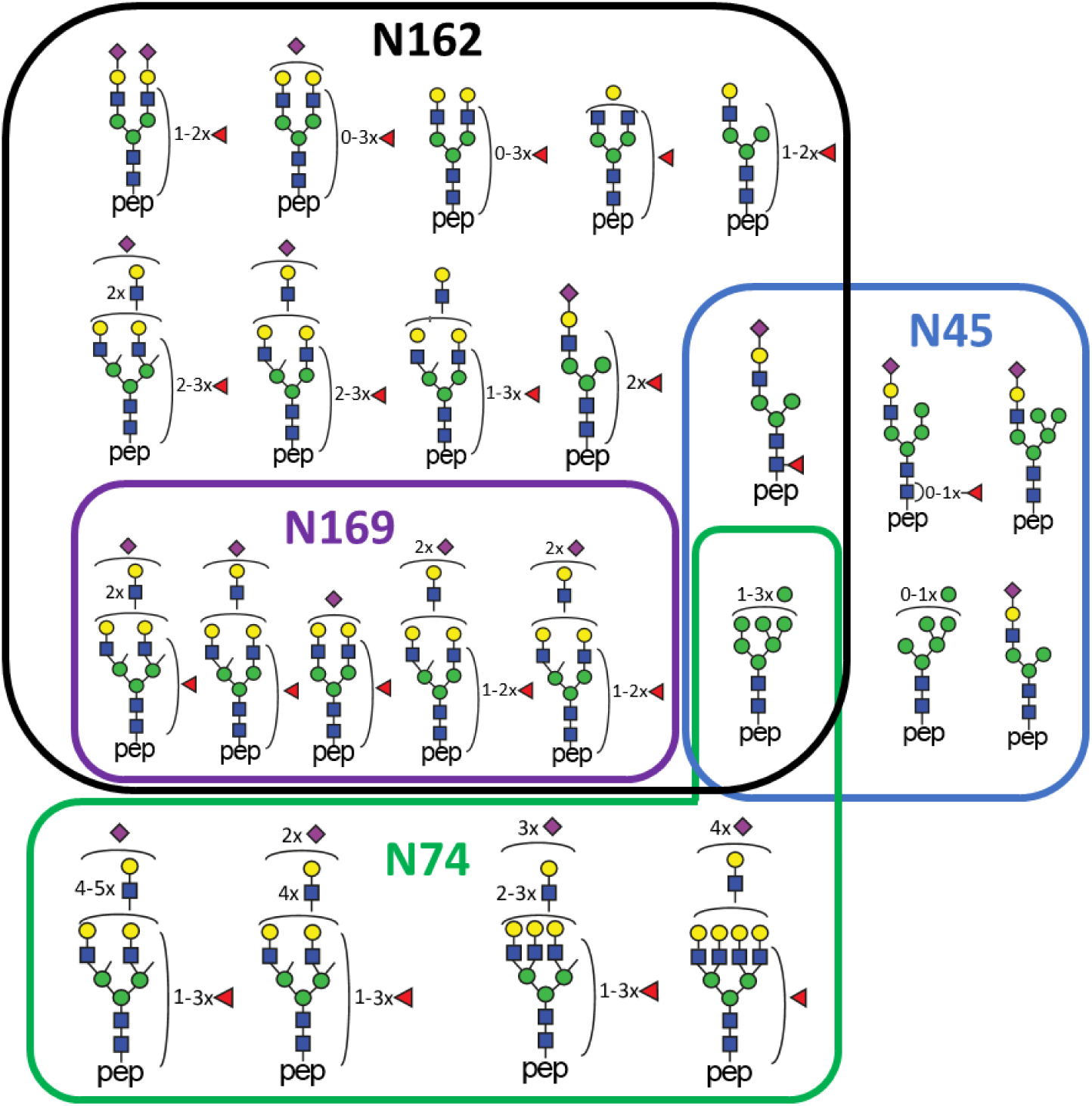
Putative structures of N-glycans identified on the four occupied N-glycosylation sites, namely 10 at N_45_, 15 at N_74_, 30 at N_162_ and 6 at N_169_. Isomeric structures co-occur and are discussed in the text. The scheme underlines the diversity, overlap and uniqueness of glycans compositions.

Two novel (N_74_, N_169_) and three already described (N_45_, N_64_, N_162_) *N*-glycosylation sites of neutrophil-derived FcγRIIIb were identified. Per site, the nature and of number the glycans differs, and, among other things reflects the extend of biosynthetic processing. The glycan heterogeneity ranges from oligomannose type glycans to highly processed complex type glycans with LacNAc extensions. Three *N*-glycosylation sites, N_45_, N_162_ and N_169_, were found to be fully occupied. Since glycosylated and non-glycosylated forms of the peptide containing site N_74_ were detected, we concluded that this site was partially occupied. As previously reported by Washburn *et al*. ^31^, the site N_64_ was not occupied and thus corresponding peptides were exclusively non-glycosylated. Molecular dynamics simulations of the highly homologous FcγRIIIa (V158 allotype) have shown intramolecular interactions between the peptide backbone residues 60 to 70 and glycans at N_45_, which may explain a preference for an unoccupied site N_64_23. Moreover, this intramolecular interaction may inhibit enzymatic *N*-glycan processing in the Golgi, providing an explanation for the observed restricted processing at site N_45_ compared to the other sites. The occupancy of site N_64_ appears to be the most prominent difference between the membrane-bound and the soluble version of the receptor IIIb (**Supplementary Table 6**). In contrast to the neutrophil-derived version, soluble FcγRIIIb has been described to display highly branched glycan structures on site N_64_ ^32^. The limited *m/z* range up to 1500 may not allow us to detect those complex structures. However, in contrast to site N_38_, the non-glycosylated peptides containing site N_64_ were detected. Furthermore, a previous study on neutrophil-bound FcγRIIIb also reported site N_64_ to be unoccupied^31^. These differences raise a lot of questions, especially concerning the glycosylation profiles of resting neutrophil-bound FcγRIIIb molecules in comparison to soluble molecules released by activated neutrophils^37^.

An example of an MS spectrum obtained for sites N_162_ and N_45_ with annotation of the major glycoforms is given in **Figure 4** and **Supplementary Figure S3**. For site N_162_ complex di- and triantennary glycans were found accompanied by a small percentage of high mannose glycans. Site N_45_ mainly showed oligomannosidic glycans with a significant fraction of hybrid and complex structures. N_74_ predominantly elaborated as di- to tera-antennary complex glycans with a small amount high mannose type glycans. N_169_ was found to exclusively carry di- and tri-antennary complex glycans.

**Figure 4.**
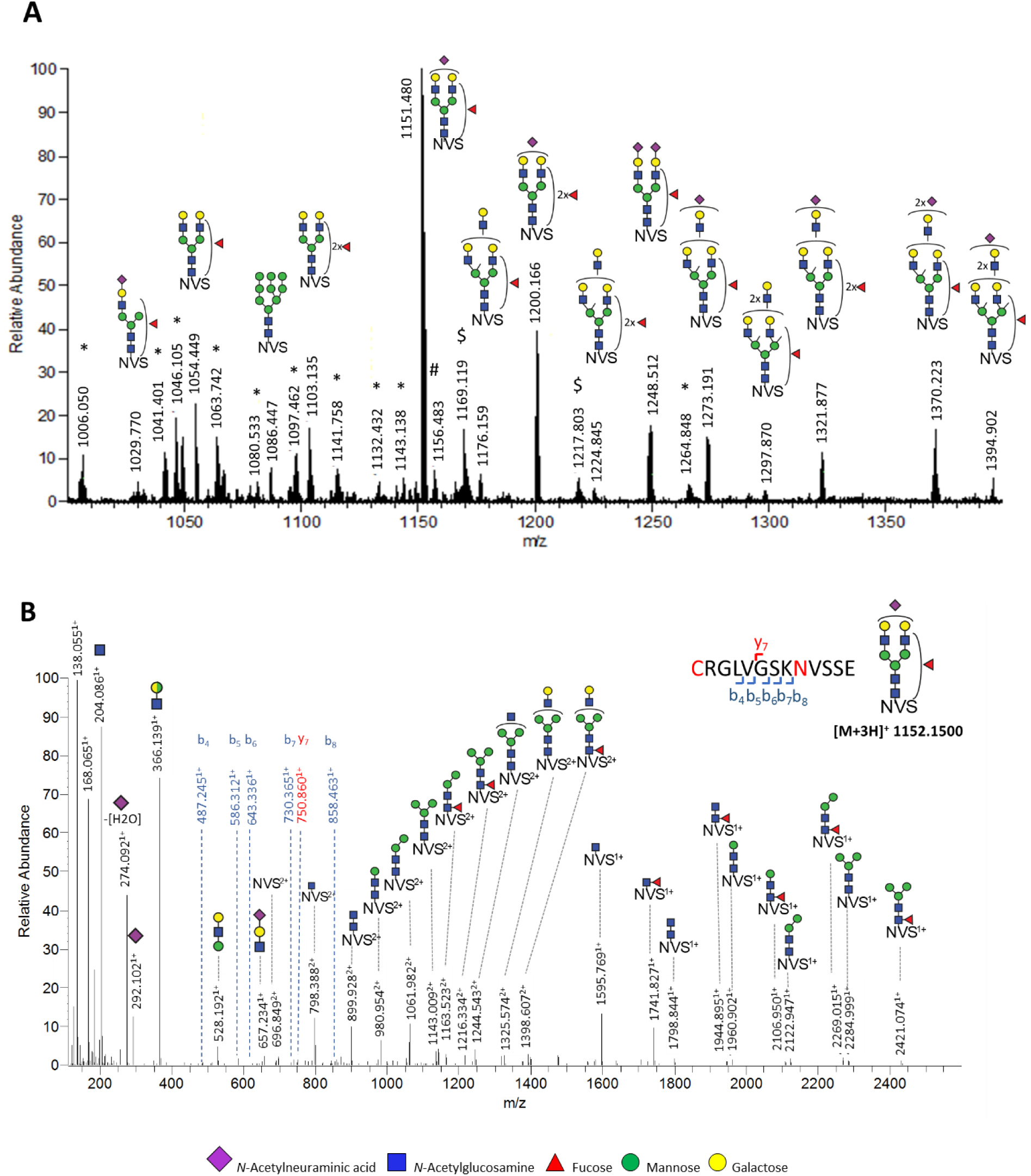
Sum spectra showing the major glycoforms of the N_162_ site. (A) MS sum spectrum (retention time 10.2 to 14.4 min) showing the major glycoforms of the N_162_ site. (B) Stepping-energy HCD MS/MS sum spectrum of the precursor ion 1152.150^3+^ [second isotopologue] of the most abundant N_162_ glycopeptide (H5N4F1S1). The MS/MS spectrum was generated for three stepping energies (32, 37, 42 V) at retention time 11.0 min. Interestingly, ammonia adducts were exclusively observed for glycopeptides carrying an oligomannose glycan, while iron adducts were detected for both oligomannose and complex structures. For more details on adduct identification see the **Supplementary Figure S3**. *: N_162_ glycopeptides with a miscleaved peptide backbone; #: unidentified glycopeptide (z=2); $: iron adducts [M+Fe^III^]^3+^. NVS: CRGLVGSK**N**_162_VSSE peptide backbone.

### Site-specific quantification of FcγIIIb *N*-glycans from human neutrophils

The 61 identified glycopeptides were targeted for relative quantification in a site-specific manner (**Figure 5**). FcγRIIIb glycosylation of the three healthy donors displayed very similar patterns in terms of composition and abundance of glycans (**Supplementary Figure S4**). Derived glycosylation traits — complexity, number of fucoses per glycan and number of sialic acids per glycan — were calculated to facilitate the comparison of the different sites (**Supplementary Figure S5)** and with other studies (**Table S7**).

**Figure 5.**
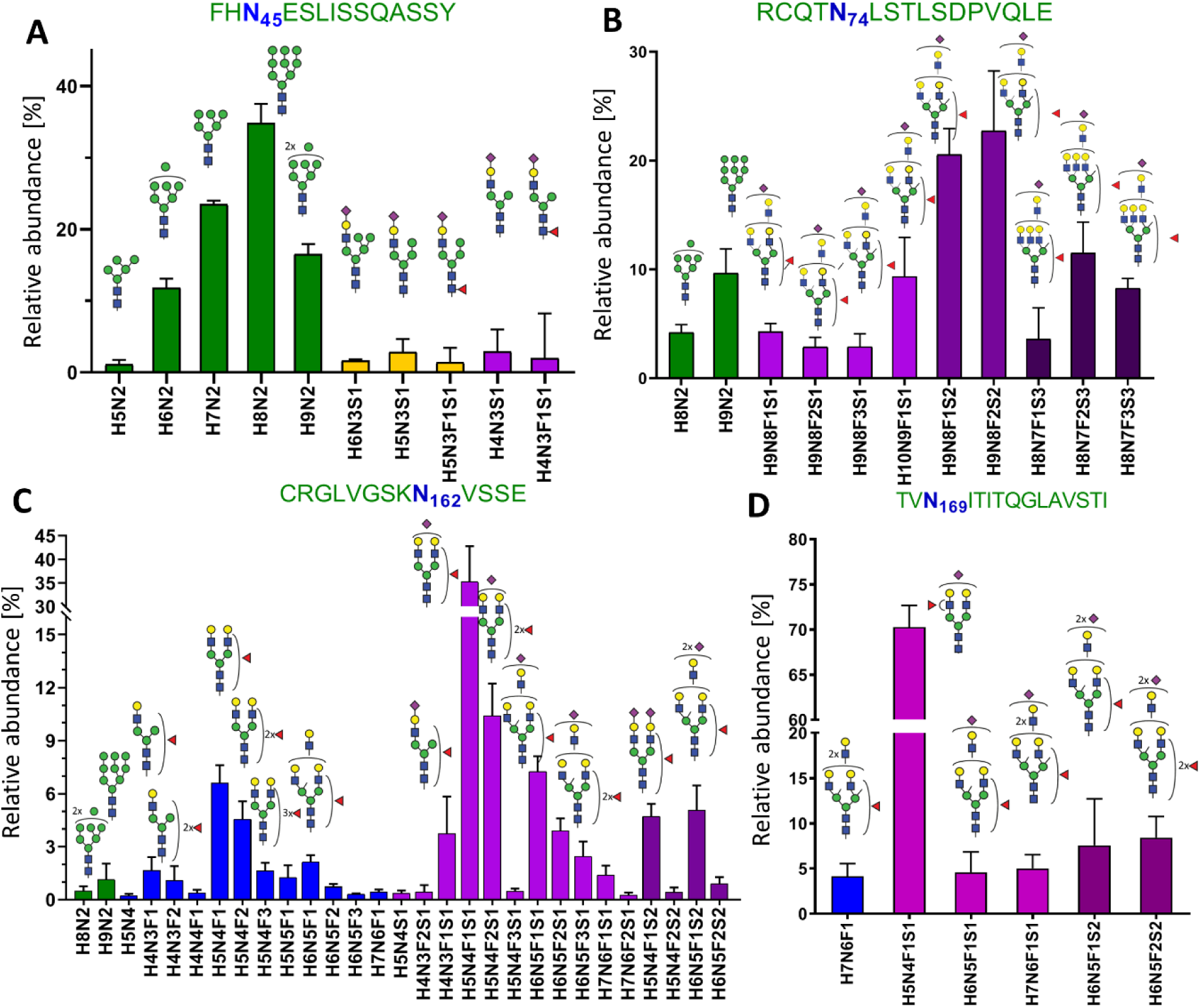
Site-specific, relative quantification of FcγRIIIb glycoforms. (A) 10 different glycan compositions occupy the glycosylation site N_45_, (B) 11 site N_74_, (C) 30 site N_162_ and (D) 6 site N_169_. Mean and SD of three donors are shown. Bar colours indicate glycan classes: Green, high mannose type; orange, hybrid type; blue, neutral complex; purple, sialylated complex, with one (light), two (medium) or three (dark) sialic acids per glycan.

The site N_45_ on FcγRIIIb is highly occupied. As shown in **Figure 5A** and **Supplementary Table S7**, approximately 86% of the glycoforms decorating this site were high mannose type glycans (M6, M7, M8, M9). The remaining glycoforms consist of 7% hybrid (M4A1G1S1, M5A1G1S1, FM4A1G1S1) and 8% complex (A1G1S1, FA1G1S1) type glycans, with and without core fucose. In contrast to serum-derived FcγRIIIb, containing only high mannose *N*-glycans^31^, the neutrophil-bound N_45_ contains a significant amount of sialylated hybrid and monoantennary complex *N*-glycans in addition.

Relative quantification of the glycosylation site N_162_ revealed 98% of sialylated, complex, mono-up to tri-antennary glycans, with some evidence of LacNAc repeats (see **Figure 5C)**. Additionally, a small percentage of high mannose glycans (2%) was observed. The derived traits revealed a high level of galactosylation and fucosylation, which indicate a high expression of both galactosyltransferases and fucosyltransferases (**Supplementary Figure 5S**). On average, 1.3 fucoses per glycan were displayed on N_162_ glycans, indicating more than one fucose residue per glycan, including core fucosylation and antennary fucosylation. Antennary fucosylation was also indicated by the B-ion observed at *m/z* 512.197^41^, corresponding to [hexose+*N*-acetylhexosamine+fucose+H]^+^. Core fucosylation was indicated by the presence of the Y-ion at *m/z* 1741.827, annotated as peptide and core fucosylated component [CRGLVGSK**N**_**162**_VSSE+*N*-acetylhexosamine+fucose+H]^+^. Some monofucosylated compositions (H5N4F1S1, H5N4F1) also showed evidence for both core and antennary fucosylated isomers. In comparison to the other glycosylation sites, N_162_ possesses the lowest amount of sialic acids per glycan (0.9), which implies poorer accessibility for sialyltransferases.

On N_74_, sialylated, fucosylated, complex *N*-glycans with multiple LacNAc extensions represent the largest group (86%) of structures (**Figure 5B**). Fragmentation of these glycans resulted in the formation of the oxonium ion of *m/z* 657.237, assigned as [*N*-acetylglucosamine-galactose-N-acetylneuraminic acid]^+^ (**Supplementary Table S8**). No evidence for the presence of oligosialylated antennae was seen in the fragments indicating that the tri-sialylated structures are at least tri-antennary. Even though N_74_ is not directly involved in antibody binding^42^, there are some speculations regarding LacNAc repetition in cell activation regulation through the modulation of receptor clustering^43,44^. Among others, this site is characterized by the highest numbers of sialic acids (2.1) and fucoses (1.5) per glycan, which implies a high accessibility for sialyltransferases and fucosyltranserases.

Site N_169_ is mainly occupied by a diantennary and monosialylated (H5N4F1S1) structure (**Figure 5D**). Interestingly, we did not observe core fucosylation as reported for serum-derived FcγRIIIb. In contrast, only B-ions indicating antennary fucosylation were observed (**Supplementary Table S5**). In general, as depicted in **Figure S5**, *N*-glycans modifying N_169_ were fully galactosylated and exhibited a moderate number of sialic acids (1.1) and fucoses (1.1) per glycan.

Overall, we confirmed the presence of antennary fucosylated glycans for three *N*-glycosylation sites, namely N_74_, N_162_, N_169_ (**Supplementary Table S5**). Among these sites, N_74_ and N_169_ were annotated as predominantly occupied by antennary fucosylated structures, since no evidence for the presence of core fucosylation was detected. In contrast, glycans on N_162_ presented a mixture of both core and antennary fucosylated isomers. In addition to complex-type species, oligomannose structures (M8, M9) complement the repertoire of complex glycans attached to N_74_ (14%) and N_162_ (2%). For N_74_, decorated by large glycans with LacNAc extension of the antennae, these oligomannose structures are being reported here for the first time. Interestingly, evidence for the presence of LacNAc units was also detected for some glycans at N_162_ (**Supplementary Table S5**). Generally, the presence of high mannose type glycoforms is not expected among highly processed glycopeptides. However, it is consistent with recent glycomics studies of FcγRIIIa^29^. The large biosynthetic gap between the high mannose type glycans and the large, (at least partially) tetraantennary glycans may indicate different subcellular fractions. It may also be linked to the accessibility of the site for mannosidases. The partial occupancy of site N_74_ suggests that accessibility may be limited early in the folding process at least for a subset of FcγRIIIb molecules. Considering the overall glycosylation pattern of FcyRIIIb in neutrophils, we estimated that nearly all glycans of FcyRIIIb were fully galactosylated, indicating a high activity of galactosyltransferases. Consequently, the partially sialylated and core fucosylated glycans suggest a moderate activity of sialyltransferases and α1,6-fucosyltransferases. Lastly, the high average number of fucose per glycan and MS/MS data for antennary fucosylation were evidence for a high activity of α1,2, α1,3 or α1,4 fucosyltransferases in the neutrophils.

### Comparison of site-specific *N*-glycosylation of FcγRIIIb

Glycosylation of sites N_45_ and N_162_ of FcγRIIIb, and the absence of it from site N_64_, has been previously reported from neutrophils of healthy donors by Washburn *et al*.^31^. The other glycosylation sites were not addressed previously. Glycosylation profiles were, both qualitatively and quantitatively, highly consistent between the study of Washburn *et al*. and ours (**Supplementary Table S6 and Table S7)**. A minor difference is the observation of high mannose *N*-glycans at site N_162_ in our study.

Recently, the site-specific *N*-glycan assignments for FcγRIIIa from human NK cells along with the experimental procedures were published^29^. We noticed major glycosylation differences in sites N_45,_ N_74_ and N_169_, clearly distinguishing the two isoforms of the FcγRIII (**Supplementary Table S6 and S7**). FcγRIIIb showed less processing than FcγRIIIa on N_45_, having mainly high mannose glycans (86%), followed by complex (8%) and hybrid glycans (7%), while FcγRIIIa expressed mainly hybrid type glycans with some complex type. The most pronounced differences were observed in the levels of antennary fucosylation. This was indicated on FcγRIIIb by an average number of fucoses per glycan higher than one, for example 1.5, 1.3 and 1.1 at sites N_74_, N_162_ and N_169_, respectively. Moreover, the fragmentation data for monofucosylated glycan compositions at site N_169_, indicated only antennary fucosylated isomers. In stark contrast, glycoforms of FcγRIIIa are reported to be almost exclusively core fucosylated. For the most abundant FcγRIIIa glycoforms, MS/MS spectra showed only glycopeptide fragments carrying core fucose. Small amounts of antennary fucosylated glycans were reported at sites N_74_ and N_162_. The difference in antennary fucosylation hints at a higher activity of α1,2, α1,3 and α1,4 fucosyltransferases in the neutrophils than in the NK cells. Among all glycosylation sites, *N*-glycans at N_162_ showed the highest similarity between the two receptor isotypes. This observation suggests that the glycosylation profiles of this functionally relevant site are conserved among FcγRIII isoforms and cells. However, sialylation at N_162_ from FcγRIIIb appeared to be lower (40%) when compared to the sialylation of FcγRIIIa from NK cells (62%). FcγRIIIb and FcγRIIIa share 97% sequence homology, but their affinity to IgG differs significantly^45^. The greater IgG binding affinity of FcγRIIIa has been attributed to a glycine residue at the position 129 in comparison to FcγRIIIb with an aspartic acid residue at position 129^46^. In the future, it might be interesting to study, whether the observed glycosylation differences have a further impact on IgG binding or another functional role.

Noticeable differences in glycosylation profiles between FcγRIII receptors may influence protein properties and function^47^. To study this functional impact, it is important to use the expressing system that assembles the revealed endogenous-like glycosylation profile. Among all five mammalian system producing differently processed *N*-glycans (HEK293, CHO, BHK, NS0)^33,48,49^, CHO cells constitute a good expression vehicle, where the highly-branched N_74_ glycans in FcγRIIIa were carrying LacNAc repetitions. However, to produce and investigate antennary fucosylation, which was observed at high levels for FcγRIIIb, the HEK293 system constitutes a better vehicle. Nevertheless, glycoengineering provides us with the possibilities to create specific FcγRIIIb structures in order to study the functional properties of different processing levels^50,51^. As shown in a mutagenesis study of the recombinant FcγRIIIa and its mouse orthologue FcγRIV, *N*-glycans at N_162_ were a critical element for antibody binding^25,52^. The present data indicate that the neutrophil-derived FcγRIIIb *N*-glycosylation profile is rather consistent between healthy individuals but significantly differs from recombinant soluble FcγRIIIb, serum-derived FcγRIIIb and NK cell-derived FcγRIIIa profiles.

## CONCLUSION

In this study, we describe a straightforward and comprehensive site-specific profiling of FcγRIIIb *N*-glycosylation with a resolution of a single donor. For the first time, all *N*-glycosylation sites were characterized in a single LC-MS/MS assay. Our approach unifies sample preparation, as 1) it uses a single proteolytic cleavage regime to characterize all sites and 2) is applicable to both FcγRIIIa and FcγRIIIb and thus potentially to many different leukocytes. It is also simpler, for example in the removal of IgG interferences and the avoidance of glycopeptide enrichment, promising more robustness. Simplicity, robustness, single donor resolution and comprehensiveness are key attributes for the use in (large-scale) clinical studies. Our method therefore presents an ideal stepping stone towards the investigation of the role of FcγRIII glycosylation for different (sub-)cellular entities in various (patho-)physiological conditions. With continued development, it has the potential to be a critical tool in the search for clinical markers of FcyRIII glycosylation.

Our analysis of the endogenous human FcyRIIIb on neutrophils of healthy donors is largely consistent with literature^31,32^. However, our improved methodology has led to several new insights. Especially, the observed differences between the plasma-derived and the neutrophil-derived FcγRIIIb demonstrate a significant biological diversity. Additionally, while site N_162_ glycosylation is highly similar, the other sites show significant variation between FcγRIIIb and FcγRIIIa glycosylation. Of note, other roles are already emerging, as exemplified for the potential impact of site N_74_ LacNAc repeats on receptor clustering ^29,43^. It is therefore prudent to equally investigate all glycosylation sites, as this is likely to provide more biological, clinical and pharmacological insights.

It would be of great interest to compare FcγRIIIb glycosylation profiles of subcellular fractions or FcγRIIIb from resting neutrophils to soluble receptor in the same donor. Additional isomers differentiation would also be desirable, but would likely need a separation method with a higher degree of isomers separation.

We believe that a throughput-optimized adaptation of the presented approach could be used in defining glycan signatures of FcγRIII for different pathophysiological conditions in various cell types or even subcellular compartments. This would reveal a yet hidden layer of regulation of antibody-mediated (auto-)immune responses. However, sensitivity should be further improved for such aims. A better understanding of glycosylation as an additional layer of regulation of FcγR activity is likely to improve the performance of antibody-based therapeutic interventions and provide clinical markers for personalized medicine in the long run.

## MATERIALS AND METHODS

### Materials

Endoproteinase Gluc-C (Staphyloccoccus aureus Protease V8) and chymotrypsin were obtained from (Worthington Biochemical Corp., Lakewood, USA). The ultrapure milli-Q deionized water (MQ) was generated using a Q-Gard 2 system (Millipore, Amsterdam, Netherlands) maintained at ≤ 18.2 MΩ. MS grade acetonitrile (ACN) was acquired from Biosolve B.V. (Valkenswaard, The Netherlands). Iodoacetamide (IAA), dithiothreitol (DTT), trizma hydrochloride, Tris(hydroxymethyl)aminomethane, Protease Inhibitor Cocktail (Set V, EDTA-Free), Nonidet P-40 substitute (NP-40), phenylmethylsulfonyl fluoride (PMSF), ethylenediaminetetraacetic acid (EDTA), NaF and glycerol were obtained from Sigma-Aldrich (Steinheim, Germany). Analytical grade formic acid (FA) and water of LC-MS grade were purchased from Fluka (Steinheim, Germany). Di-sodium hydrogen phosphate dihydrate (Na_2_HPO_4_·2 H_2_O), potassium dihydrogen phosphate (KH_2_PO_4_), and NaCl were obtained from Merck (Darmstadt, Germany).CARIN lysis buffer (pH 8.0) was prepared in-house with 20 mM Tris-HCl, 137 mM NaCl, 10 mM EDTA, 0.1 M NaF, 1% NP-40 and 10% glycerol. Phosphate-buffered saline (PBS, 0.035 M, pH 7.6) was prepared in-house with 5.7 g/L of Na_2_HPO_4_, 2 H_2_O, 0.5 g/L of KH_2_PO_4_, and 8.5 g/L of NaCl. Coomassie staining was prepared in-house according to Candiano *et al*. ^54^ using Coomassie Blue G-250 (Sigma-Aldrich). FcγRIII were immunoprecipitated from the neutrophil cell lysate using a mouse anti-CD16 monoclonal IgG2a antibody (Ref M9087, Clone CLB-FcR gran/1, 5D2, Sanquin, Amsterdam, The Netherlands). Prior to usage, antibodies were labeled with biotin. At first, antibodies were buffer exchanged from Tris buffer to PBS buffer using the Zeba spin protocol (ThermoFisher Scientific, Rockford, IL, USA), as amine-containing buffers may interfere with biotinylation. Subsequently, the Z-link™ Sulfo-NHS-Biotinylation protocol (ThermoFisher Scientific) was followed. The level of biotin incorporation was measured with a HABA Assay (ThermoFisher Scientific).

### Neutrophil cell isolation and FcγRIIIb purification

Neutrophils were isolated from whole blood of three healthy volunteers as described previously^55^. Initially, the blood was anticoagulated with EDTA. Mononuclear leukocytes and platelets were then removed by centrifugation using a Ficoll gradient (Ficoll-Paque PLUS, GE Healthcare, Freiburg, Germany). Erythrocytes were subsequently lysed with isotonic NH_4_Cl solution at 4°C resulting in a cellular fraction of neutrophils, at a concentration of 50 × 10^6^ cells/mL (**Supplementary Table S1**). Isolated neutrophils were then washed two times with 1 mL of cold PBS. Afterwards, they were lysed in CARIN lysis buffer with protease inhibitors for 10 min on ice (Protease Inhibitor Cocktail and PMSF). The cell lysate was centrifuged at 13000 × *g* for 15 min at 4°C. The supernatant was then transferred and the cellular debris pellet was discarded. Based on the total protein content measured with a NanoDrop™ 2000 spectrophotometer (ThermoFisher Scientific, Rockford, IL, USA), 100 µg or 500 µg of neutrophil proteins was incubated, while rotating, with 5 µg or 25 µg of biotinylated antibodies in CARIN lysis buffer overnight at 4°C. 10 µL of High Capacity Streptavidin Agarose Resin beads (ThermoFisher Scientific) were washed twice in 1 mL CARIN buffer and incubated with the pre-formed FcγRIII-anti-CD16 immune complexes for 1 hour at 4°C and rotating. Thereafter, the beads were centrifuged for 2 min at 2500 × *g* and the supernatant discarded. The beads were then washed four times with 1 mL CARIN buffer. Finally, FcγRIII was eluted from the beads with two times 150 µL of 200 mM FA. The eluates were then dried by vacuum centrifugation at 60°C for 2 hours.

### SDS-PAGE and western blotting

The immunoprecipitation of FcγRIIIb from 100 µg of total neutrophil proteins was evaluated by sodium dodecyl-sulfate-polyacrylamide gel electrophoresis (SDS-PAGE). The dried samples were resuspended in 20 µL of non-reducing loading buffer (NuPAGE™ LDS Sample Buffer (4x), ThermoFisher Scientific, Germany) and incubated at 95°C for 5 min. The samples as well as protein standards (PageRuler Prestained Protein Ladder, ThermoFisher Scientific) were then separated on a 4-12% Bis-Tris gel (NuPAGE™, ThermoFisher Scientific) in a 4-Morpholinepropanesulfonic acid (MOPS) SDS running buffer (NuPAGE™, ThermoFisher Scientific). The migration was performed at 200 V constant voltage for 50 min. The obtained gels were stained using Commassie blue. The presence of CD16 was confirmed by western blotting, using an anti-CD16 mouse monoclonal IgG1 conjugated to horseradish peroxidase (DJ130c, sc-20052 HRP, Lot #D2617, Santa Cruz Biotech). At first, proteins were transferred from the gel onto a polyvinylidene difluoride (PVDF) membrane using an iBlot™2 Gel Transfer Device (ThermoFisher Scientific) at 20 V for 6 min. The membrane was then washed three times for 5 min with 1X Tris-Buffered Saline Tween 20 (TBST) and blocked with 5% milk (Elk, Campina, Amersfoort, The Netherlands) in TBST for 30 min. It was washed again thrice in TBST for 10 min before being incubated overnight at 4°C in 5% milk TBST with the anti-CD16 antibody diluted 1000 times. Thereafter, the membrane was washed three times for 10 min in TBST and then treated with enhanced chemiluminescent substrate (Pierce™ ECL Western Blotting Substrate, ThermoFisher Scientific). Both gels and western blots were visualized using a ChemiDoc MP Imaging system (Biorad, Hercules, USA), under Trans-UV for gels and chemiluminescence detection for membranes.

### In-gel proteolytic digest and LC-MS glycopeptide analysis

In the case of the in-gel LC-MS workflow the, upscaled amount of 500 µg of total neutrophil proteins was used for immunoprecipitation. The subsequent SDS-PAGE was performed at 200 V constant voltage for only 15 min. The obtained gels were silver stained (SilverQuest™ Staining Kit, Invitrogen) and the protein of interest was cut out from the gel. Excised bands were subjected to reduction with 10 mM DTT, followed by alkylation with 50 mM IAA^56^. The in-gel digestion steps with endoproteinase GluC and chymotrypsin were performed by a Proteineer DP digestion robot (Bruker, Bremen, Germany)^33^. The liquid chromatography-tandem mass sprectometry (LC-MS/MS) analysis was performed on a nanoLC-MS system composed of an Easy nLC 1000 gradient HPLC system (Thermo, Bremen, Germany) and a LUMOS mass spectrometer (Thermo). Prior to sample injection, the peptides and glycopeptides extracted from the bands were lyophilized and dissolved in solvent A (3% ACN/95% water containing 0.1% FA (v/v)). The samples were then loaded onto an in-house packed C18 precolumn (100 μm × 15 mm; Reprosil-Pur C18-AQ 3 μm, Dr. Maisch, Ammerbuch, Germany) and separated on a homemade analytical nanoLC column (30 cm × 75 μm; Reprosil-Pur C18-AQ 3 μm). The digested (glyco-)peptides were eluted using a linear gradient from 10 to 40% solvent B (80% ACN/20% water containing 0.1% FA (v/v)) over 20 min. The nanoLC column was drawn to a tip of ∼5 μm and acted as the electrospray needle of the MS source. The Orbitrap Fusion LUMOS mass spectrometer was operated in data-dependent MS/MS (top-20 mode). The master spectra (MS1) were acquired within a mass range of *m/z* of 400–1500 with an AGC target value of 1 × 10^4^ for an accumulation time of maximum 50 ms. The resolution setting for MS1 scans was 12 × 10^4^. Dynamic exclusion duration was 10 s with a single repeat count, and charge states in the range 1–5 were included for MS/MS. The resolution of MS/MS scans was 3 × 10^4^ with an AGC target of 5 × 10^4^ with maximum fill time of 60 ms MS/MS spectra were generated from precursors isolated with the quadrupole with an isolation width of 1.2 Da. The acquisition of collision-induced dissociation (CID) spectra was performed for the precursor and recorded at 35 V. In addition, the higher-energy collisional dissociation (HCD) data acquisition was triggered by the presence of the *N*-acetylhexosamine (HexNAc) oxonium ion at *m/z* 204.087. Upon the glycan diagnostic ion recognition, an HCD collision stepping-energies of 32, 37 and 42 V were applied and three additional data-depended MS/MS scans of the same precursor were executed. The following parameters were set: AGC target of 5 × 10^5^ with a fill time of max 200 ms. The presented results on protein identification, coverage and purity are from a standard data-dependent HCD run (without exclusive *m/z* 204.087 triggering). For the identification of glycopeptides in these runs, MS/MS spectra containing the specific HexNAc oxonium ion at m/z 204.087 (HexNAc, [C_8_H_14_NO_5_]^+^) were extracted from the raw data and written to an .mgf file using in-house routines. This filtering step also ensured to include only spectra containing the HexNAc oxonium ion with a strong signal (among top 30 peaks).

### Identification and quantification of site-specific glycosylation

Initially, the acquired LC-MS/MS data were processed by Byonic software package (Protein Metrics, Cupertino, CA v3.2-38)^57^. Byonic’s Glycopeptide Search was used for automatic peptide and glycopeptide detection by searching fragmentation data (MS/MS) against a database containing an extensive human proteome combined with a pre-defined glycan composition list (**Supplementary Table S8**). For the search, the digestion specificity parameter was set as non-specific (slowest), allowing both specific and miscleaved peptides to be included in the search. The glycosylation was set as a common modification and other anticipated modifications were set based upon prevalence: Glycan modification/ +[glycan composition] Da @ NGlycan | common1; carbamidomethyl/ +57.021 Da @C | fixed; oxidation/ +15.995 Da @M | common2; acetyl/ +42.011 Da @Protein *N*-term | rare1. The remining parameters were set as shown in **Supplementary Table S9**. Further, the list of FcγRIII glycoforms determined by Byonic was manually verified and completed by the identification of the remaining glycoforms in Xcalibur (Thermo). **Supplementary Table S10** gives an overview of the level (Byonic, manual MS/MS, manual MS) at which the individual glycoforms were identified. Regarding the manual identification, MS1 sum spectra were generated around the retention times where glycopeptides had been assigned. This was done per combination of unique peptide backbone and number of sialic acids, the two features which are mainly influencing the retention time. Some of sialic acid variants were inferred, improving the identification of multisialylated glycan compositions. The sum spectra were then manually searched for expected monosaccharide differences. Annotation of MS1 spectra was based on precursor mass with a tolerance of ± 0.05 Th and confirmed with the manual interpretation of the MS/MS spectra of precursor-ions. Manual MS/MS interpretation was based on, firstly the clear presence of an *m/z* corresponding to a peptide or peptide+HexNAc fragment ion of a previously identified glycosylated sequence and secondly the presence of a dominating pattern of oxonium ions. During the manual interpretation, differences in (glyco-)peptide sequences between the FcγRIIIb allotypes, NA1 and NA2 (**Supplementary Table S4**), were taken into consideration.

For automatic alignment and integration of LC-MS data, the in-house software LacyTools (Version 1.1.0-alpha) was used as described previously^58^. The following settings were applied: sum spectrum resolution of 100, extraction mass window of 0.07 Th, extraction time window of 15 s, percentage of the theoretical isotopic pattern of 95%. After extraction, analytes were curated from the identification list, if the average mass error was outside ±20 ppm and the isotopic pattern deviated more than 20% from the theoretical one. The intensities of the iron adduct ([M+Fe^III^]^3+^) and ammonia adduct ([M+2H+NH_4_]^3+^) signals were significant. Hence, the relative quantification was performed on extracted areas of protonated, iron adduct and ammonia adduct peaks. Total area normalization per glycosylation site was used for relative quantitation.

## ACKNOWLEDGEMENTS

We thank Gillian Dekkers for expressing the recombinant receptors used in the initial development of the method. This research was supported by the European Union (Glycosylation Signatures for Precision Medicine project, GlySign, Grant No. 722095), and the Netherlands Organization for Scientific Research (NWO) via Vernieuwingsimpuls Veni (Project No. 722.016.008) and Medium Investment (to P.A.V.; 91116004; partly financed by ZonMw) grants.

## DATA AVAILABILITY

The raw mass spectrometric data files that support the findings of this article are available from the corresponding author upon request.

## AUTHOR CONTRIBUTIONS

I.W., T.S., A.H.R. and G.M.C.J. carried out all experiments, partially assisted by D.F.. E.L.G. provided the neutrophil samples. T.S. optimized the immunoprecipitation procedure with support from E.L.G. I.W. performed the immunoprecipitation of the donor samples. G.M.C.J. performed and optimized the in-gel digestion. A.H.R. performed and advised on the LC-MS/MS measurements and database searches. Y.M. contributed computational tools. I.W. and D.F. processed and analyzed the data. D.F., M.W., G.V. and P.A.V. designed and supervised the study. I.W., D.F. and T.S. wrote the manuscript with significant contribution from all authors.

