## Supplementary Information for "Site-specific glycosylation mapping of Fc gamma receptor IIIb from neutrophils of health donors"

***Both authors contributed equally**

### SUPPLEMENTARY FIGURES

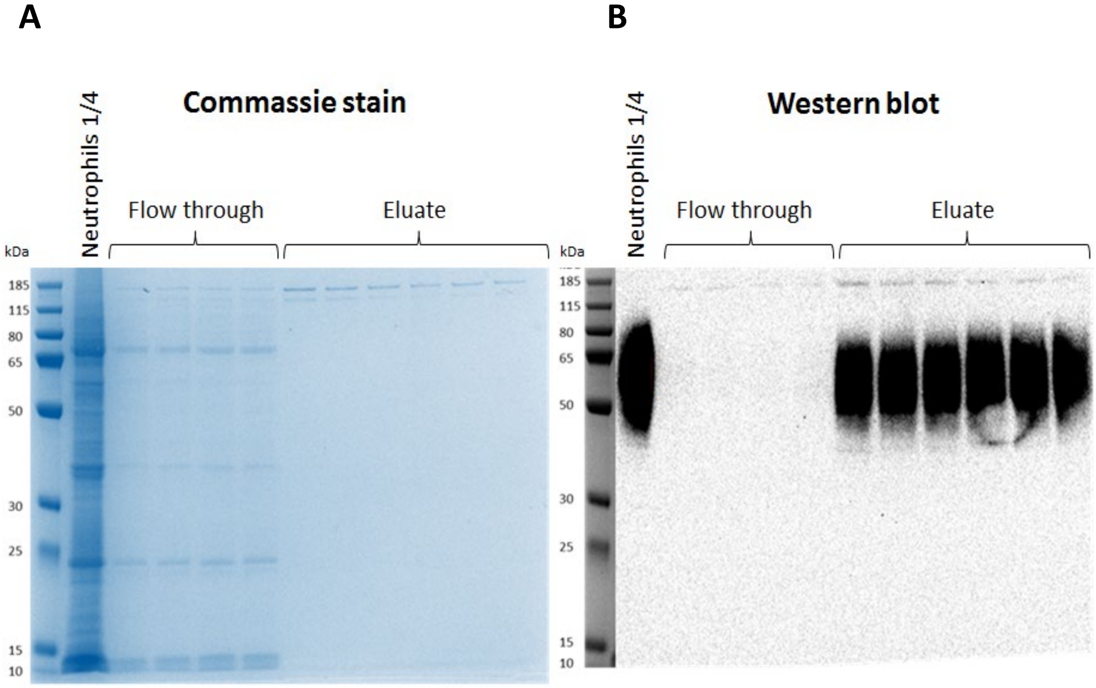


**Figure S1.** Non-reducing SDS-PAGE and western blot of human neutrophil lysate before and after FcγRIII immunoprecipitation. Protein contaminations and FcγRIIIb were detected with (A) commassie staining and (B) western blot, respectively. The neutrophils lanes represent ~25 µg of total protein content from the neutrophil cell lysate from donor 2. In this experiment, the immunoprecipitation was done from 100 µg and the eluate was split between the two gels (50 µg per gel). The flow-through lanes show the unbound diluted fraction of the immunoprecipitation while the eluate lanes present the purified FcγRIIIb protein. As expected, FcγRIIIb exhibits an elongated band from 50 to 80 kDa, visible after western blotting. Despite high dilution, the flow-through mirrors the major bands from the total cell lysate. In contrast, the eluate exhibits none of those bands and thus any interfering proteins of 50-80 kDa above the limit of detection (LOD; 25 ng protein). Two bands around 150 kDa appear in the flow-through and eluate. They are likely derived from the capturing antibody, as an in-solution proteolytic cleavage of the eluate revealed immunoglobulin G glycopeptide masses (data not shown). Whatever the identity of the contamination, it is the main reason to prefer an in-gel proteolytic cleavage. At an input of 16 million neutrophils and an estimated 100,000 to 300,000 receptors per cell with a molecular weight of 50 to 80 kDa, the expected maximum yield should be around 130 to 630 ng[1]. In the eluate, the FcγRIIIb could not be detected by Coomassie blue staining (LOD ca. 25 ng for well-defined bands). This indicated a low protein recovery. However, a strong enrichment of FcγRIIIb from neutrophil lysate was achieved (Supplementary Table S2).

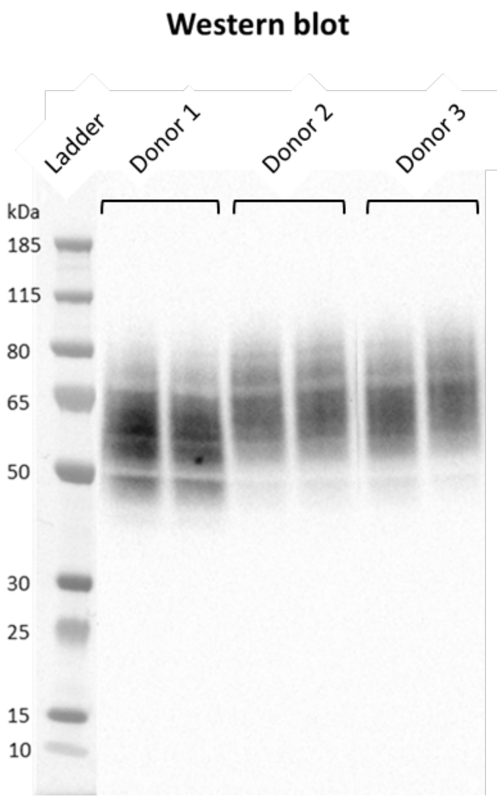


**Figure S2.** The immunoblot shows FcγRIIIb immunoprecipitated from the lysate of neutrophils. Lanes contain material from donor 1 (NA1/NA2), donor 2 (NA2/NA2) and donor 3 (NA2/NA2) in duplicate. Size markers are indicated on the left. FcγRIIIb showed a smear from 50 to 80 kDa indicating the heterogeneity of the *N*-glycosylation pattern of the receptor. Differential electrophoretic mobility of the receptor from NA1/NA2-heterozygous and NA2-homozygous is likely mainly due to NA polymorphism and distinct number of N-glycosylation sites[2]. Larger proteoforms attributed to NA2, may not be visible in the first donor, because the protein expression is skewed towards NA1.

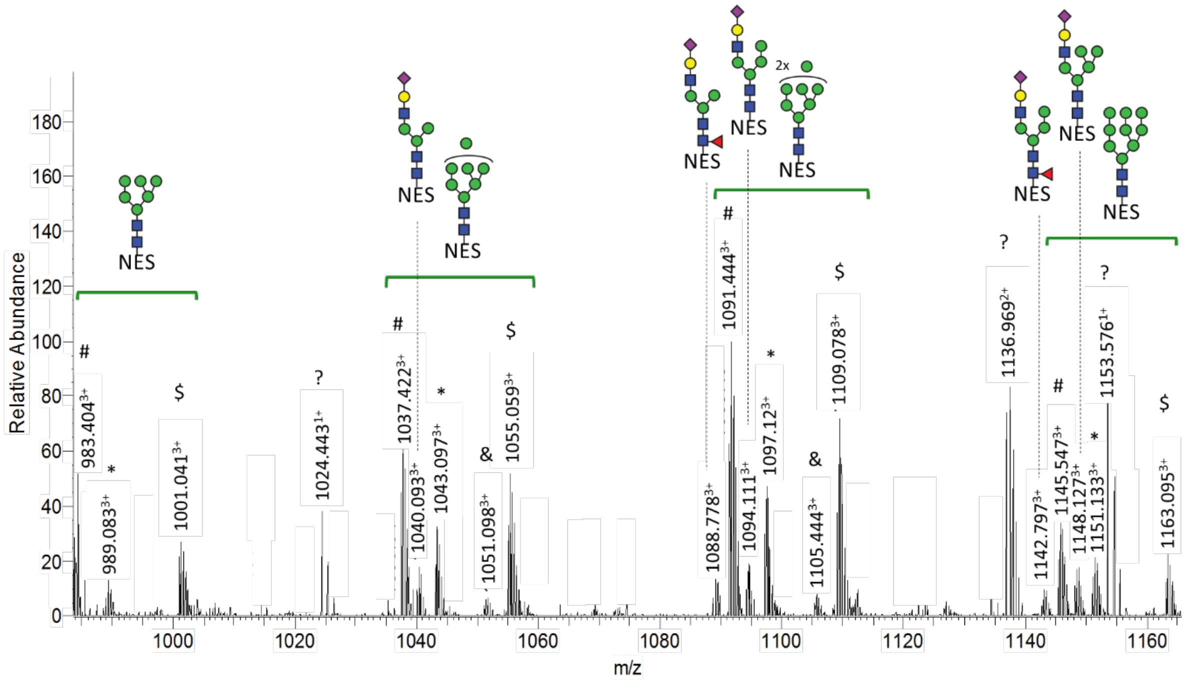


**Figure S3**. Sum spectra showing the different ions observed for the $N_{45}$ glycoforms of FcγRIIIb. The mass spectrum was generated for donor 2 within the following retention time range: 17.4-19.4 min. NES: FH**N₄₅**ESLISSQASSY peptide backbone. The MS spectra revealed satellite peaks for some triply-charged protonated species with a difference of 55.934 Da and 18.033 Da. These were identified by accurate mass (< ± 5 ppm) as iron ($[M+Fe^{III}]^{3+}$) and ammonia ($[M+2H+NH_4]^{3+}$) adducts, respectively. They also presented the same retention time as the respective protonated glycopeptide. Additionally, the presence of the ammonia adducts was confirmed by MS/MS spectra, as the comparison of the b- and y-ion series provided evidence for the same peptide backbone without additional modification. #: the triply protonated $N_{45}$ glycopeptides $[M+3H]^{3+}$. *: the triply charged ions of protonated $N_{45}$ glycopeptides with ammonia adducts $[M+2H+NH_4]^{3+}$. $: $N_{45}$ glycopeptides charged by iron ions $[M+Fe^{III}]^{3+}$. &: misscleaved glycopeptide. ?: unidentified glycopeptide

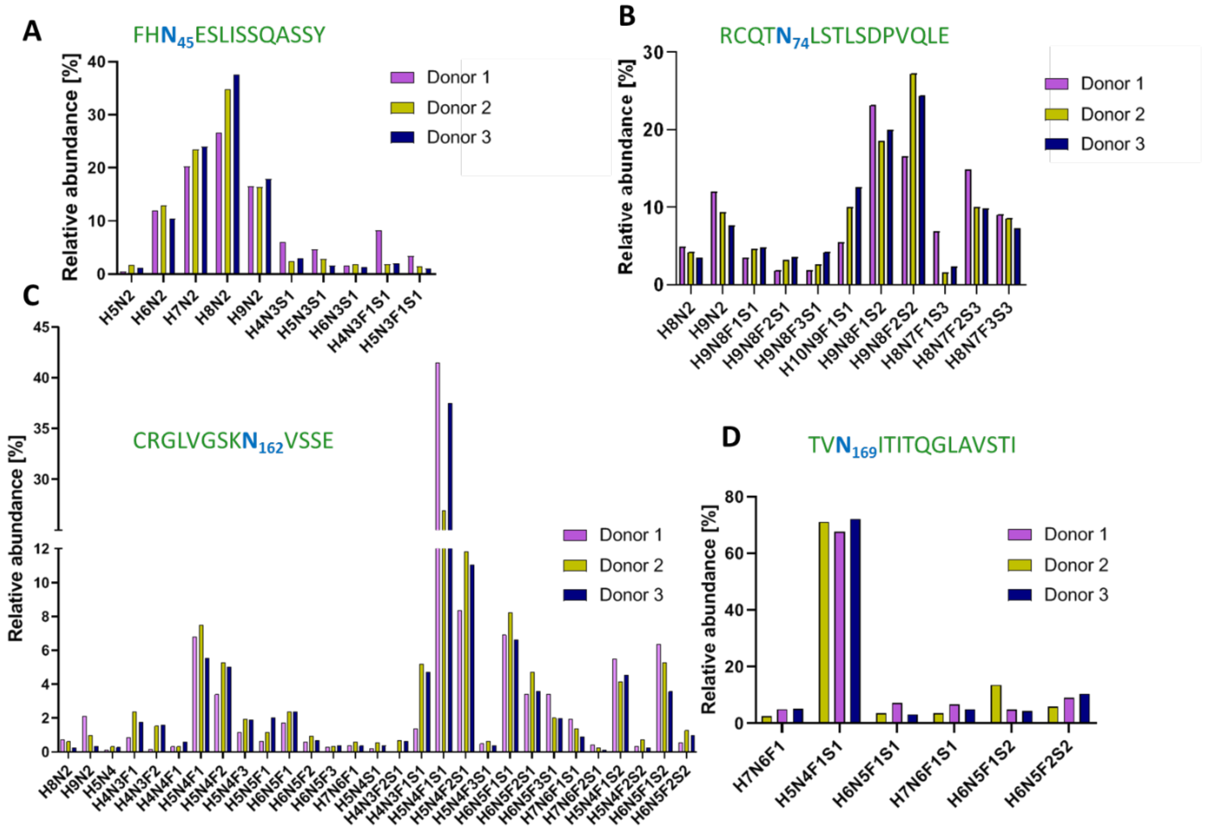


**Figure S4.** Site-specific relative quantification of FcγRIIIb glycoforms. (A) 10 different glycan compositions occupy glycosylation site $N_{45}$, (B) 11 site $N_{74}$, (C) 30 site $N_{162}$ and (D) 6 site $N_{169}$.

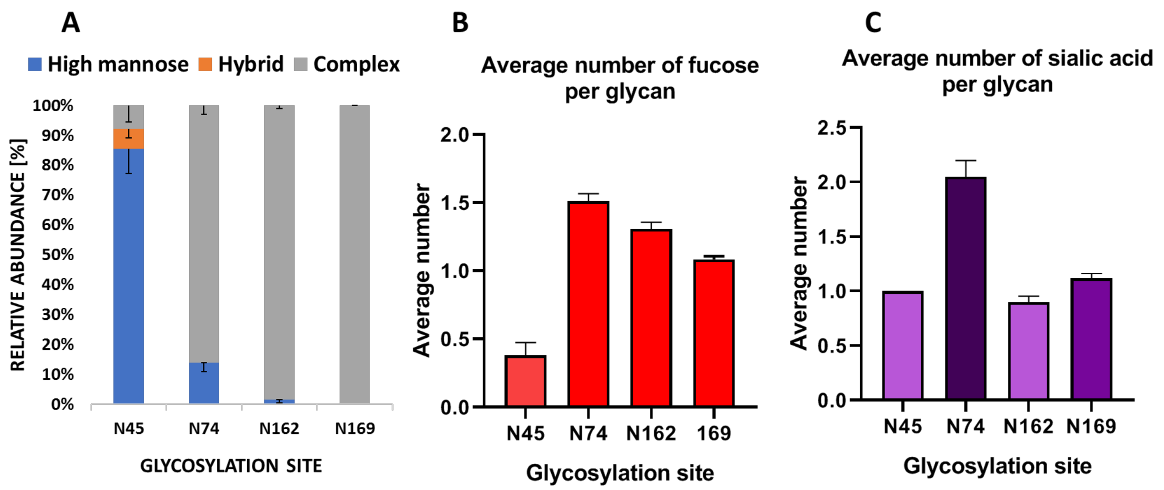


**Figure S5.** Various glycosylation parameters were calculated to evaluate the glycosylation profile of FcγRIIIb. Displayed is the average and standard deviation per glycosylation site of the three donors regarding several derived glycosylation traits: (A) the relative abundance of the three basic N-glycans types; (B) the average number of fucoses per glycan; (C) the average number of sialic acids per glycan. For detailed calculation of the presented traits refer to Supplementary Table S9.

### SUPPLEMENTARY TABLES

**Table S1.** Information on healthy donors and separated neutrophils cells used for FcγRIIIb isolation.

| Donor information | Donor 1 | Donor 2 | Donor 3 |
|---|---|---|---|
| Sex | female | male | female |
| Cell type | neutrophils | neutrophils | neutrophils |
| Cell count [cell/mL] | $50x10^6$ | $50x10^6$ | $50x10^6$ |
| Total protein con. [mg/mL] | 1.61 | 1.72 | 1.51 |
| Amount of cells for isolation [cell/mL] | $16x10^6$ | $15x10^6$ | $17x10^6$ |
| Volume of lysate for isolation [µL] | 310 | 291 | 331 |
| Amount of total protein for isolation [µg] | 500 | 500 | 500 |

**Table S2.** Byonic output of critical parameters reflecting the quality of the peptide-spectrum matches (PSMs) of identified FcγRIIIb for each donor. The list of parameters includes: protein p-value (the likelihood of the PSMs to be the identified protein). # unique peptides (total number of PSMs). coverage % (percent of the protein sequence covered by PSMs).

<table>
<tr><th>Donor</th><th>Protein Identified</th><th>Protein p-value (Log base 10)</th><th># unique peptides</th><th>Ranking place</th><th>Coverage</th></tr>
<tr><td>1</td><td rowspan="3">Low affinity immunoglobulin gamma Fc region receptor III-B UNIPROT: O75015 (FCG3B_HUMAN)</td><td>362.01</td><td>131</td><td>1</td><td>77.8 %</td></tr>
<tr><td>2</td><td>268.28</td><td>127</td><td>1</td><td>80.3 %</td></tr>
<tr><td>3</td><td>250.22</td><td>114</td><td>1</td><td>71.2 %</td></tr>
</table>

**Table S3.** Characterization of predominant human FcγRIIIb (UniProtKB - O75015) site-specific peptides and glycopeptides generated from Endoproteinase Glu-C and chemotrypsin cleavage. C, carboxymethyl cysteine

| Site | Peptide sequence | Glycosylation | *m/z* peptide [M+H] | *m/z* peptide [M+GlcNAc] | RT [min] |
|---|---|---|---|---|---|
| Asn 38 | missing peptide | NA | NA | NA | NA |
| Asn 45 | FHN(45)ES(47)LISSQASSY | + | 1569.708 | 1772.787 | 17 - 23 |
| Asn 64 | FIDAATVN(64)DSGEY | - | 1401.617 | 1604.696 | 19 |
| Asn 74 | RCQTN(74)LSTLSDPVQLE | + | 1860.901 | 2063.980 | 20-28 |
| | RCQTN(74)LSTLSDPVQLE | - | 1860.901 | - | 25 |
| Asn 162 | CRGLVGSKN(162)VSSE | + | 1392.679 | 1595.758 | 10 - 14 |
| Asn 169 | TVN(169)ITITQGLAV | + | 1229.706 | 1432.779 | 28 - 32 |

**Table S4.** Allelic peptide and glycopeptide sequences containing N45 and N64/D64 used for polymorphic variant assessment. *Uniprot identifiers.

| FcγRIII variant | $N_{45}$ peptide | | $N_{64}$ peptide |
|---|---|---|---|
| NA1 *O75015 (FCG3B*01) | $FHN_{45}EN(47)LISSQASSY$ | | $FIDAATVD_{64}DSGEY$ |
| NA2 *O75015 (FCG3B*02) | $FH\mathbf{N}_{45}ES(47)LISSQASSY$ | | $FIDAATV\mathbf{N}_{64}DSGEY$ |
| FcγRIIIa *P08637 | $FH\mathbf{N}_{45}ES(47)LISSQASSY$ | | $FIDAATVD_{64}DSGEY$ |
| DONOR | RATIO ALLOTYPIC | | ALLOTYPE |
| | NES/(NES+NEN) | NDS/(DDS+NDS) | |
| Donor 1 | 0.2 | 0.26 | NA1/NA2 |
| Donor 2 | 0.99 | 0.98 | NA2/NA2 |
| Donor 3 | 0.99 | 0.98 | NA2/NA2 |

**Table S6.** Site-specific derived glycosylation traits. Average number of sialic acids and fucoses per glycan of endogenous neutrophil-derived FcγRIIIb, soluble FcγRIIIb and NK-derived FcγRIIIa. *the corresponding glycopeptides were not detected in this study **potential glycosylation site was not occupied by oligosaccharides, only corresponding peptide observed. For detailed calculations of the traits, see **Table S10**.

| Sites | Derived traits | FcγRIIIb – our study | sFcγRIIIb[3] (human serum) | FcγRIIIb[4] (human neutrophils) | FcγRIIIa[5] (human NK cells) |
|---|---|---|---|---|---|
| $N_{38}$ | average number of sialic acids | -* | 3.3 | - * | 4.0 |
| | average number of fucoses | | 2.2 | | 1.0 |
| $N_{64}$ | average number of sialic acids | -** | 2.5 | - ** | - |
| | average number of fucoses | | 1.3 | | |
| $N_{45}$ | average number of sialic acids | 1.0 | 0.0 | 1.0 | 1.2 |
| | average number of fucoses | 0.5 | 0.0 | 0.5 | 0.5 |
| $N_{74}$ | average number of sialic acids | 2.1 | 3.7 | - * | 4.0 |
| | average number of fucoses | 1.5 | 2.0 | | 1.1 |
| $N_{162}$ | average number of sialic acids | 0.9 | 1.1 | 1.1 | 1.3 |
| | average number of fucoses | 1.3 | 1.2 | 1.2 | 1.1 |
| $N_{169}$ | average number of sialic acids | 1.1 | 1.5 | - * | 1.5 |
| | average number of fucoses | 1.1 | 1.2 | | 1.0 |

**Table S7.** The glycosylation profiles of endogenous neutrophil-derived FcγRIIIb, soluble FcγRIIIb and NK-derived FcγRIIIa. The different glycoforms are displayed as defined derived traits (high mannose, complex, hybrid). *the corresponding glycopeptides were not detected in this study **potential glycosylation site was not occupied by oligosaccharides, only corresponding peptide observed.

| Sites | FcγRIIIb – our study (human neutrophils) | sFcγRIIIb[3] (human serum) | FcγRIIIb[4] (human neutrophils) | FcγRIIIa[6] (human natural killer cells) |
|---|---|---|---|---|
| $N_{38}$ | -* | Complex (LacNAc repeats) > 99% | - * | Complex (LacNAc repeats) > 99% |
| $N_{45}$ | Complex (monoantennary) 8%<br>Hybrid 7%<br>High mannose 86% | High mannose >99% | Complex (monoantennary) 16%<br>Hybrid < 30%<br>High mannose 54% | Complex (di-, tri-, tetra antennary) 22%<br>Hybrid 71%<br>High mannose 2% |
| $N_{64}$ | -** (not glycosylated) | Complex (LacNAc repeats) > 99% | - ** (not glycosylated) | - |
| $N_{74}$ | Complex (LacNAc repeats) 86%<br>High mannose 14% | Complex (LacNAc repeats) > 99 % | - * | Complex (LacNAc repeats) > 99% |
| $N_{162}$ | Complex (mono-, di- , tri antennary) 98%<br>High mannose 2% | Complex (di-, tri-, tetraantennary) > 99% | Complex (di-, tri antennary) > 99% | Complex (di-, tri-antennary) 66%<br>Hybrid 22%<br>High mannose 12% |
| $N_{169}$ | Complex (di-, tri-, tetrantennary) > 99% | Complex (di-, tri-, tetrantennary) > 99% | - * | Complex (di-, tri-, tetrantennary) > 99% |

**Table S9.** The search parameters used for initial analysis using Byonic (Protein Metrics, Cupertino, CA v3.2-38).

| Search parameter | Value |
|---|---|
| Protein database | UniProt human protein database |
| Glycan database | User defined, see Table S2 |
| Precursor mass tolerance | 5 ppm |
| Fragmentation type | QTOF/HCD |
| Parent tolerance | 0.005 *m/z* |
| Fragment mass tolerance | 20 ppm |
| Recalibration (lock mass) | none |
| Maximum precursor mass | 10,000 |
| Precursor and mass charge assignments | compute from MS1 |
| Max # of precursor per scan | 2 |
| Smoothing width | 0.01 *m/z* |

**Table S10.** Site-specific derived trait calculation. The compositions are placeholders for their respective relative intensities.

| Site | NUMBER OF SIALIC ACIDS<br>Average number of sialic acids per glycan on complex glycans of FcyRIIIb |
|---|---|
| $N_{45}$ | (H4N3S1 + H4N3F1S1) / (H4N3S1 + H4N3F1S1) |
| $N_{74}$ | (H9N8F1S1 + H9N8F2S1 + H9N8F3S1 + H10N9F1S1) + 2*(H9N8F1S2 +H9N8F2S2) + 3*(H8N7F1S3+H8N7F2S3+H8N7F3S3)) / (H9N8F1S1 + H9N8F2S1 + H9N8F3S1 + H10N9F1S1 + H9N8F1S2 + H9N8F2S2 + H8N7F1S3 + H8N7F2S3 + H8N7F3S3) |
| $N_{162}$ | (H5N4S1 + H4N3F2S1 + H4N3F1S1 + H5N4F1S1 + H5N4F2S1 + H5N4F3S1 + H6N5F1S1 + H6N5F2S1 + H6N5F3S1 + H7N6F1S1 + H7N6F2S1) + 2*(H5N4F1S2 + H5N4F2S2 + H6N5F1S2 + H6N5F2S2)) / (H5N4 + H4N3F1 + H4N3F2 + H4N4F1 + H5N4F1 + H5N4F2 + H5N4F3 + H5N5F1 + H6N5F1 + H6N5F2 + H6N5F3 + H7N6F1 + H5N4S1 + H4N3F2S1 + H4N3F1S1 + H5N4F1S1 + H5N4F2S1 + H5N4F3S1 + H6N5F1S1 + H6N5F2S1 + H6N5F3S1 + H7N6F1S1 + H7N6F2S1 + H5N4F1S2 + H5N4F2S2 + H6N5F1S2 + H6N5F2S2) |
| $N_{169}$ | (H5N4F1S1 + H6N5F1S1 + 2*(H6N5F1S2 + H6N5F2S2) + H7N6F1S1) / (H5N4F1S1 + H6N5F1S1 + H6N5F1S2 + H6N5F2S2 + H7N6F1S1 + H7N6F1) |
| Site | NUMBER OF FUCOSES<br>Average number of fucoses per glycan on complex glycans of FcyRIIIb |
| $N_{45}$ | (H4N3F1S1) / (H4N3S1 + H4N3F1S1) |
| $N_{74}$ | (H9N8F1S1 + H10N9F1S1 + H9N8F1S2 + H8N7F1S3) + 2*(H9N8F2S1 + H9N8F2S2+H8N7F2S3) + 3*(H9N8F3S1+H8N7F3S3) / ( H9N8F1S1 + H9N8F2S1 + H9N8F3S1 + H10N9F1S1 + H9N8F1S2 + H9N8F2S2 + H8N7F1S3 + H8N7F2S3 + H8N7F3S3) |
| $N_{162}$ | (H4N3F1 + H4N4F1 + H5N4F1 + H5N5F1 + H6N5F1 + H7N6F1 + H4N3F1S1 + H5N4F1S1 + H6N5F1S1 + H7N6F1S1 + H5N4F1S2 + H6N5F1S2 + 2*(H4N3F2 + H5N4F2 + H6N5F2 + H4N3F2S1 + H5N4F2S1 + H6N5F2S1 + H7N6F2S1 + H5N4F2S2 + H6N5F2S2) + 3*(H5N4F3 + H6N5F3 + H5N3F3S1 + H6N5F3S1) / (H5N4 + H4N3F1 + H4N3F2 + H4N4F1 + H5N4F1 + H5N4F2 + H5N4F3 + H5N5F1 + H6N5F1 + H6N5F2 + H6N5F3 + H7N6F1 + H5N4S1 + H4N3F2S1 + H4N3F1S1 + H5N4F1S1 + H5N4F2S1 + H5N4F3S1 + H6N5F1S1 + H6N5F2S1 + H6N5F3S1 + H7N6F1S1 + H7N6F2S1 + H5N4F1S2 + H5N4F2S2 + H6N5F1S2 + H6N5F2S2) |
| $N_{169}$ | (H7N6F1 + H7N6F1S1 + H5N4F1S1 + H6N5F1S1 + H6N5F1S2 + 2*H6N5F2S2) / (H5N4F1S1 + H6N5F1S1 + H6N5F1S2 + H6N5F2S2 + H7N6F1S1 + H7N6F1) |
| Site | HIGH MANNOSE<br>Fraction of high mannose glycans |
| $N_{45}$ | (H5N2 + H6N2 + H7N2 + H8N2 + H9N2) / (H5N2 + H6N2 + H7N2 + H8N2 + H9N2 + H4N3S1 + H5N3S1 + H6N3S1 + H4N3F1S1 + H5N3F1S1) |
| $N_{74}$ | (H8N2 + H9N2) / (H8N2 + H9N2 + H9N8F1S1 + H9N8F2S1 + H9N8F3S1 + H10N9F1S1 + H9N8F1S2 + H9N8F2S2 + H8N7F1S3 + H8N7F2S3 + H8N7F3S3) |
| $N_{162}$ | (H8N2 + H9N2) / (H8N2 + H9N2 + H5N4 + H4N3F1 + H4N3F2 + H4N4F1 + H5N4F1 + H5N4F2 + H5N4F3 + H5N5F1 + H6N5F1 + H6N5F2 + H6N5F3 + H7N6F1 + H5N4S1 + H4N3F2S1 + H4N3F1S1 + H5N4F1S1 + H5N4F2S1 + H5N4F3S1 + H6N5F1S1 + H6N5F2S1 + H6N5F3S1 + H7N6F1S1 + H7N6F2S1 + H5N4F1S2 + H5N4F2S2 + H6N5F1S2 + H6N5F2S2) |
| $N_{169}$ | - |
| Site | HYBRID<br>Fraction of hybrid glycans |
| $N_{45}$ | (H5N3S1 + H6N3S1 + H5N3F1S1) / (H5N2 + H6N2 + H7N2 + H8N2 + H9N2 + H4N3S1 + H5N3S1 + H6N3S1 + H4N3F1S1 + H5N3F1S1) |
| $N_{74}$ | - |
| $N_{162}$ | - |
| $N_{169}$ | - |
| Site | COMPLEX<br>Fraction of complex glycans |
| $N_{45}$ | (H4N3S1 + H4N3F1S1) / (H5N2 + H6N2 + H7N2 + H8N2 + H9N2 + H4N3S1 + H5N3S1 + H6N3S1 + H4N3F1S1 + H5N3F1S1) |
| $N_{74}$ | (H9N8F1S1 + H9N8F2S1 + H9N8F3S1 + H10N9F1S1 + H9N8F1S2 + H9N8F2S2 + H8N7F1S3 + H8N7F2S3 + H8N7F3S3) / (H8N2 + H9N2 + H9N8F1S1 + H9N8F2S1 + H9N8F3S1 + H10N9F1S1 + H9N8F1S2 + H9N8F2S2 + H8N7F1S3 + H8N7F2S3 + H8N7F3S3) |
| $N_{162}$ | (H5N4 + H4N3F1 + H4N3F2 + H4N4F1 + H5N4F1 + H5N4F2 + H5N4F3 + H5N5F1 + H6N5F1 + H6N5F2 + H6N5F3 + H7N6F1 + H5N4S1 + H4N3F2S1 + H4N3F1S1 + H5N4F1S1 + H5N4F2S1 + H5N4F3S1 + H6N5F1S1 + H6N5F2S1 + H6N5F3S1 + H7N6F1S1 + H7N6F2S1 + H5N4F1S2 + H5N4F2S2 + H6N5F1S2 + H6N5F2S2) / (H8N2 + H9N2 + H5N4 + H4N3F1 + H4N3F2 + H4N4F1 + H5N4F1 + H5N4F2 + H5N4F3 + H5N5F1 + H6N5F1 + H6N5F2 + H6N5F3 + H7N6F1 + H5N4S1 + H4N3F2S1 + H4N3F1S1 + H5N4F1S1 + H5N4F2S1 + H5N4F3S1 + H6N5F1S1 + H6N5F2S1 + H6N5F3S1 + H7N6F1S1 + H7N6F2S1 + H5N4F1S2 + H5N4F2S2 + H6N5F1S2 + H6N5F2S2) |
| $N_{169}$ | (H7N6F1 + H5N4F1S1 + H6N5F1S1 + H7N6F1S1 + H6N5F1S2 + H6N5F2S2) / (H7N6F1 + H5N4F1S1 + H6N5F1S1 + H7N6F1S1 + H6N5F1S2 + H6N5F2S2) |

1 Huizinga, T. W., Roos, D. & von dem Borne, A. E. Neutrophil Fc-gamma receptors: a two-way bridge in the immune system. *Blood* **75**, 1211-1214 (1990).
2 Huizinga, T. W. *et al.* Soluble Fc gamma receptor III in human plasma originates from release by neutrophils. *J Clin Invest* **86**, 416-423, doi:10.1172/JCI114727 (1990).
3 Yagi, H. *et al.* Site-specific N-glycosylation analysis of soluble Fcgamma receptor IIIb in human serum. *Sci Rep* **8**, 2719, doi:10.1038/s41598-018-21145-y (2018).
4 Washburn, N. *et al.* Characterization of Endogenous Human FcgammaRIII by Mass Spectrometry Reveals Site, Allele and Sequence Specific Glycosylation. *Mol Cell Proteomics* **18**, 534-545, doi:10.1074/mcp.RA118.001142 (2019).
5 Patel, K. R., Nott, J. D. & Barb, A. W. Primary Human Natural Killer Cells Retain Proinflammatory IgG1 at the Cell Surface and Express CD16a Glycoforms with Donor-dependent Variability. *Mol Cell Proteomics* **18**, 2178-2190, doi:10.1074/mcp.RA119.001607 (2019).
6 Patel, K. R., Nott, J. D. & Barb, A. W. Primary human natural killer cells retain proinflammatory IgG1 at the cell surface and express CD16a glycoforms with donor-dependent variability. *Mol Cell Proteomics*, doi:10.1074/mcp.RA119.001607 (2019).
